# A zombie *LIF* gene in elephants is up-regulated by TP53 to induce apoptosis in response to DNA damage

**DOI:** 10.1101/187922

**Authors:** Juan Manuel Vazquez, Michael Sulak, Sravanthi Chigurupati, Vincent J. Lynch

## Abstract

Large bodied organisms have more cells that can potentially turn cancerous than small-bodied organisms with fewer cells, imposing an increased risk of developing cancer. This expectation predicts a positive correlation between body size and cancer risk, however, there is no correlation between body size and cancer risk across species (‘Peto’s Paradox’). Here we show that elephants and their extinct relatives (Proboscideans) may have resolved Peto’s Paradox in part through re-functionalizing a *leukemia inhibitory factor* pseudogene (*LIF6*) with pro-apoptotic functions. The *LIF6* gene is transcriptionally up-regulated by TP53 in response to DNA damage, and translocates to the mitochondria where it induces apoptosis. Phylogenetic analyses of living and extinct Proboscidean *LIF6* genes indicates its TP53 response element evolved coincident with the evolution of large body sizes in the Proboscidean stem-lineage. These results suggest that re-functionalizing of a pro-apoptotic LIF pseudogene may have been permissive (though not sufficient) for the evolution of large body sizes in Proboscideans.

## Introduction

The risk of developing cancer places severe constraints on the evolution of large body sizes and long lifespans in animals. If all cells have a similar risk of malignant transformation and equivalent cancer suppression mechanisms, organism with many cells should have a higher risk of developing cancer than organisms with fewer cells. Similarly organisms with long lifespans have more time to accumulate cancer-causing mutations than organisms with shorter lifespans and therefore should also be at an increased risk of developing cancer, a risk that is compounded in large-bodied, long-lived organisms (Cairns, 1975; Caulin and Maley, 2011; Doll, 1971; Peto, 2015; Peto, 1975). Consistent with these expectations, there is a strong positive correlation between body size and cancer incidence *within* species. Larger dog breeds, for example, have higher rates of cancer than smaller breeds (Dobson, 2013) and human cancer incidence increases with increasing adult height for numerous cancer types (Green et al., 2011). In stark contrast, there are no correlations between body size or lifespan and cancer risk *between* species (Abegglen et al., 2015); this lack of correlation is often referred to as ‘Peto’s Paradox’ (Caulin and Maley, 2011; Leroi et al., 2003; Peto, 1975).

While the ultimate resolution to Peto’s paradox is that large bodied and/or long-lived species evolved enhanced cancer protection mechanisms, identifying and characterizing those mechanisms is essential for elucidating how enhanced cancer resistance and thus large bodies and long lifespans evolved. Numerous and diverse mechanisms have been proposed to resolve Peto’s paradox (Caulin and Maley, 2011; Dang, 2015; Katzourakis et al., 2014; Leroi et al., 2003; Maciak and Michalak, 2015; Nagy et al., 2007; Nunney, 1999; Takemoto et al., 2016), but discovering those mechanisms has been challenging because the ideal study system is one in which a large, long-lived species is deeply nested within a clade of smaller, short-lived species - all of which have sequenced genomes. Unfortunately, few lineages fit this pattern. Furthermore while comparative genomics can identify genetic changes that are phylogenetically associated the evolution of enhanced cancer protection, determining which of those genetic changes are causally related to cancer biology through traditional reverse and forward genetics approaches are not realistic for large species such as whales and elephants. Thus we must use other methods to demonstrate causality.

Among the most parsimonious mechanisms to resolve Peto’s paradox are a reduced number of oncogenes and/or an increased number of tumor suppressor genes (Caulin and Maley, 2011; Leroi et al., 2003; Nunney, 1999), but even these relatively simple scenarios are complicated by transcriptional complexity and context dependence. The multifunctional interleukin-6 class cytokine *leukemia inhibitory factor* (*LIF*), for example, can function as either a tumor suppressor or an oncogene depending on the context. Classically LIF functions as an extracellular cytokine by binding the LIF receptor (LIFR) complex, which activates downstream PI3K/AKT, JAK/STAT3, and TGFβ signaling pathways. The *LIF* gene encodes at least three transcripts, *LIF-D, LIF-M*, and *LIF-T*, which contain alternative first exons spliced to common second and third exons (Haines et al., 1999; Hisaka et al., 2004; Rathjen et al., 1990; Voyle et al., 1999). Remarkably while the LIF-D and LIF-M isoforms are secreted proteins that interact with the LIF receptor (Rathjen et al., 1990; Voyle et al., 1999), the LIF-T isoform lacks the propeptide sequence and is an exclusively intracellular protein (Haines et al., 1999; Voyle et al., 1999) that induces caspase-dependent apoptosis through an unknown mechanism (Haines et al., 2000).

Here we show that the genomes of Paenungulates (elephant, hyrax, and manatee) contain numerous duplicate *LIF* pseudogenes, at least one (*LIF6*) of which is expressed in elephant cells and is up-regulated by TP53 in response to DNA damage. LIF6 encodes a separation of function isoform structurally similar to LIF-T that induces apoptosis when overexpressed in multiple cell types and is required for the elephant-specific enhanced cell death in response to DNA-damage. These results suggest that the origin of a zombie *LIF* gene (a reanimated pseudogene that kills cells when expressed) may have contributed to the evolution of enhanced cancer resistance in the elephant lineage and thus the evolution large body sizes and long lifespans.

## Results

### Repeated segmental duplications increased *LIF* copy number in Paenungulates

We characterized *LIF* copy number in 53 mammalian genomes, including large, long-lived mammals such as the African elephant (*Loxodonta africana*), Bowhead (*Balaena mysticetus*) and Minke (*Balaenoptera acutorostrata scammoni*) whales, as well as small, long-lived mammals such bats and the naked mole rat. We found that most Mammalian genomes encoded a single *LIF* gene, however, the manatee (*Trichechus manatus*), rock hyrax (*Procavia capensis*), and African elephant genomes contained 7-11 additional copies of *LIF* (**Figure 1**). None of the duplicate *LIF* genes includes the 5’-UTR, coding exon 1, or a paired low complexity (CGAG)n/CT-rich repeat common to the canonical *LIF* genes in elephant, hyrax, manatee, tenrec, and armadillo (**Figure 2A**). Most of the duplicates include complex transposable element insertions composed of tandem tRNA-Asn-AAC/AFROSINE and AFROSINE3/tRNA-RTE/MIRc elements within introns one and two (**Figure 2A**). Fine mapping of the duplicate ends by reciprocal best BLAT indicates that there is no region of homology upstream of the tRNA-Asn-AAC/AFROSINE elements for duplicates that include exon 2, whereas duplicate *LIF* genes that lack exon 2 have ~150-300bp regions of homology just upstream of the paired AFROSINE3/tRNA-RTE/MIRc elements in intron 2. The *LIF* encoding loci in the hyrax and manatee genomes have not been assembled into large-scale scaffolds, but the African elephant *LIF* loci are located within a 3.5Mb block of chromosome 25 (loxAfr4).

**Figure 1.**
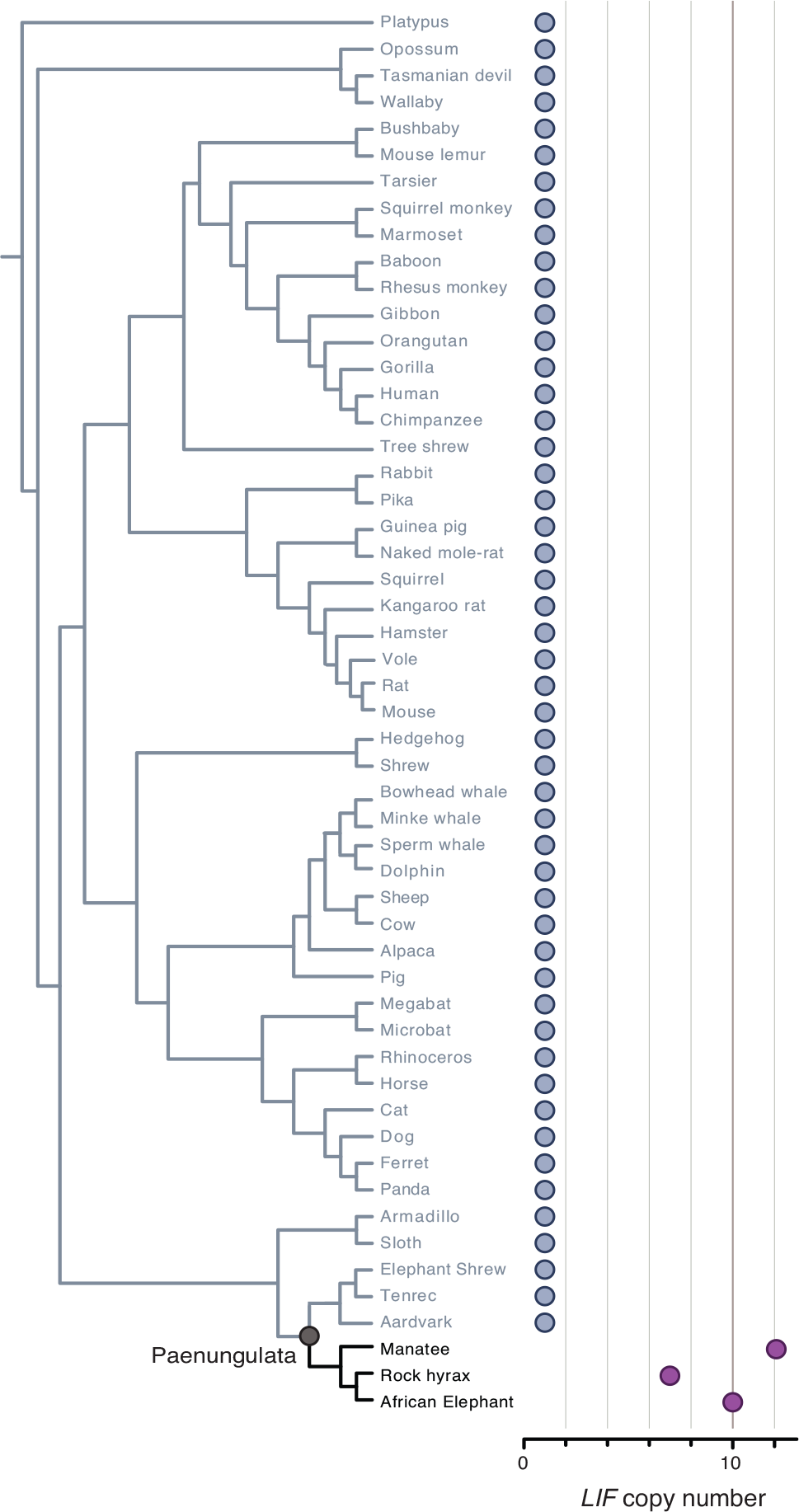
Expansion of *LIF* copy number in Paenungulata. *LIF*copy number in mammalian genomes. Clade names are shown for lineages in which the genome encodes more than one *LIF* gene or pseudogene.

**Figure 2.**
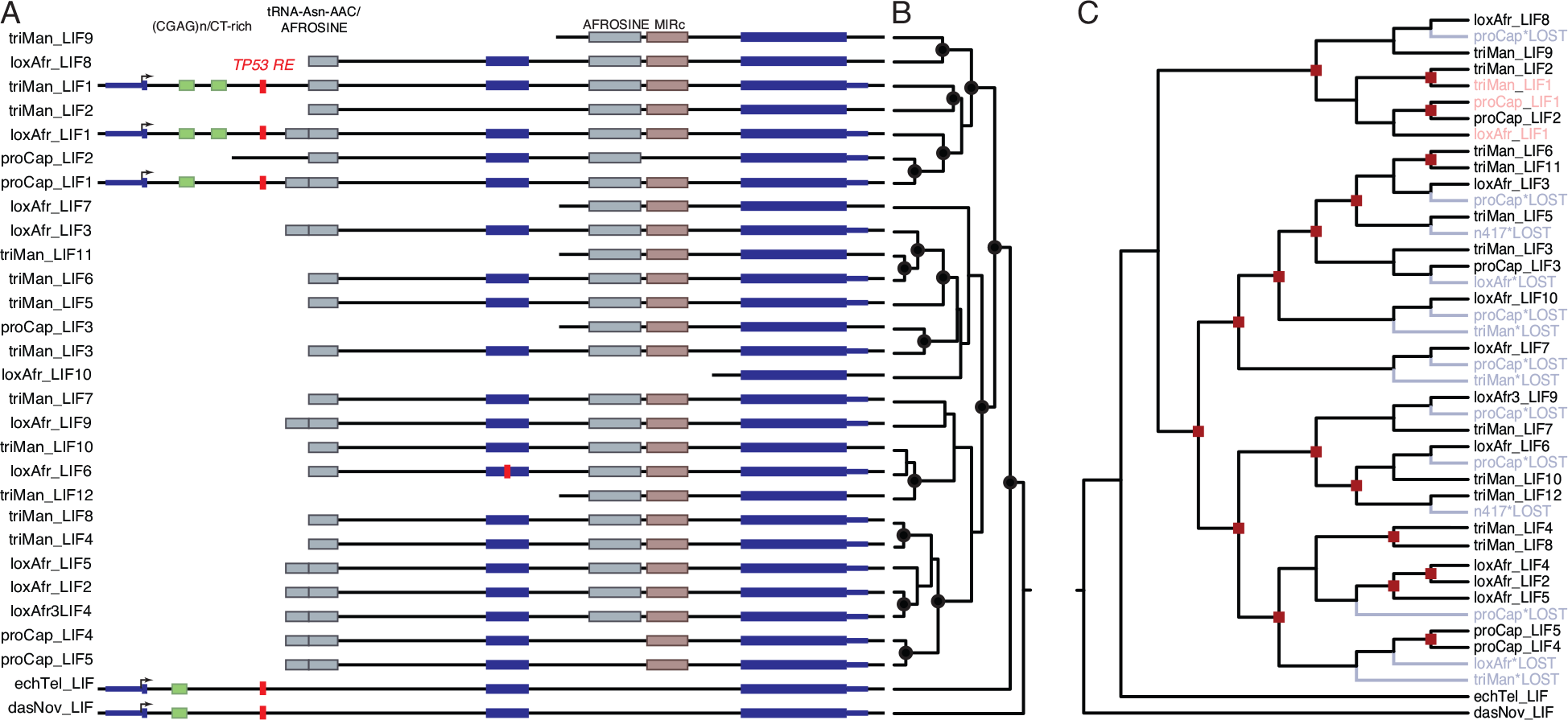
*LIF* copy number increased through segmental duplications. (A) Organization of the *LIF* loci in African elephant (loxAfr), hyrax (ProCap), and manatee (triMan), tenrec (echTel), and armadillo (dasNov) genomes. The location of homologous transposable elements around *LIF* genes and TP53 transcription factor binding sites are shown. (B) *LIF* gene tree, nodes with Bayesian Posterior Probabilities (BPP) > 0.9 are indicated with black circles. (C) Reconciled *LIF* gene trees African elephant (loxAfr), hyrax (ProCap), and manatee (triMan). Duplication events are indicated with red squares, gene loss events are indicated with in blue and noted with ‘*LOST’. Cannonical *LIF* genes (*LIF1*) are shown in red.

*LIF* duplicates may result from independent duplication events in the elephant, hyrax, and manatee lineages, ancestral duplications that occurred in the Paenungulate stem-lineage followed by lineage-specific duplication and loss events, or some combination of these processes. We used Bayesian phylogenetic methods to reconstruct the *LIF* gene tree and gene tree reconciliation to reconstruct the pattern of *LIF* duplication and loss events in Paenungulates. Consistent with a combination of ancestral and lineage-specific duplications, our phylogenetic analyses of Paenungulate *LIF* genes identified well-supported clades containing loci from multiple species as well as clades containing loci from only a single species (**Figure 2B**). The reconciled tree identified 17 duplication and 14 loss events (**Figure 2C**). These data indicate that the additional *LIF* genes result from repeated rounds of segmental duplication, perhaps mediated by recombination between repeat elements.

### Duplicate *LIF* genes are structurally similar to the LIF-T

Barring transcription initiation from cryptic upstream sites encoding in frame start codons, all duplicate *LIF* genes encode N-terminally truncated variants that are missing exon 1, lack the propeptide sequence, and are similar in primary structures to LIF-T (**Figure 3A**). While some duplicates lack the N-terminal LIFR interaction site (**Figure 3A**), all include the leucine/isoleucine repeat required for inducing apoptosis (**Figure 3A**) (Haines et al., 2000). Crucial residues that mediate the interaction between LIF and LIFR (**Figure 3B**) (Hudson et al., 1996; Huyton et al., 2007) are relatively well conserved in duplicate LIF proteins, as are specific leucine/isoleucine residues that are required for the pro-apoptotic functions of LIF-T (**Figure 3C**)(Haines et al., 2000). Haines et al. (2000) suggested that the leucine/isoleucine residues of LIF-T are located on a single face of helix B, and may form an amphipathic α-helix. Similar to LIF-T, leucine/isoleucine residues of duplicate LIF proteins are located on a single face of helix B (**Figure 3D**). These data suggest that at least some of the structural features that mediate LIF functions, in particular the pro-apoptotic function(s) of LIF-T, are conserved in duplicate LIFs.

**Figure 3.**
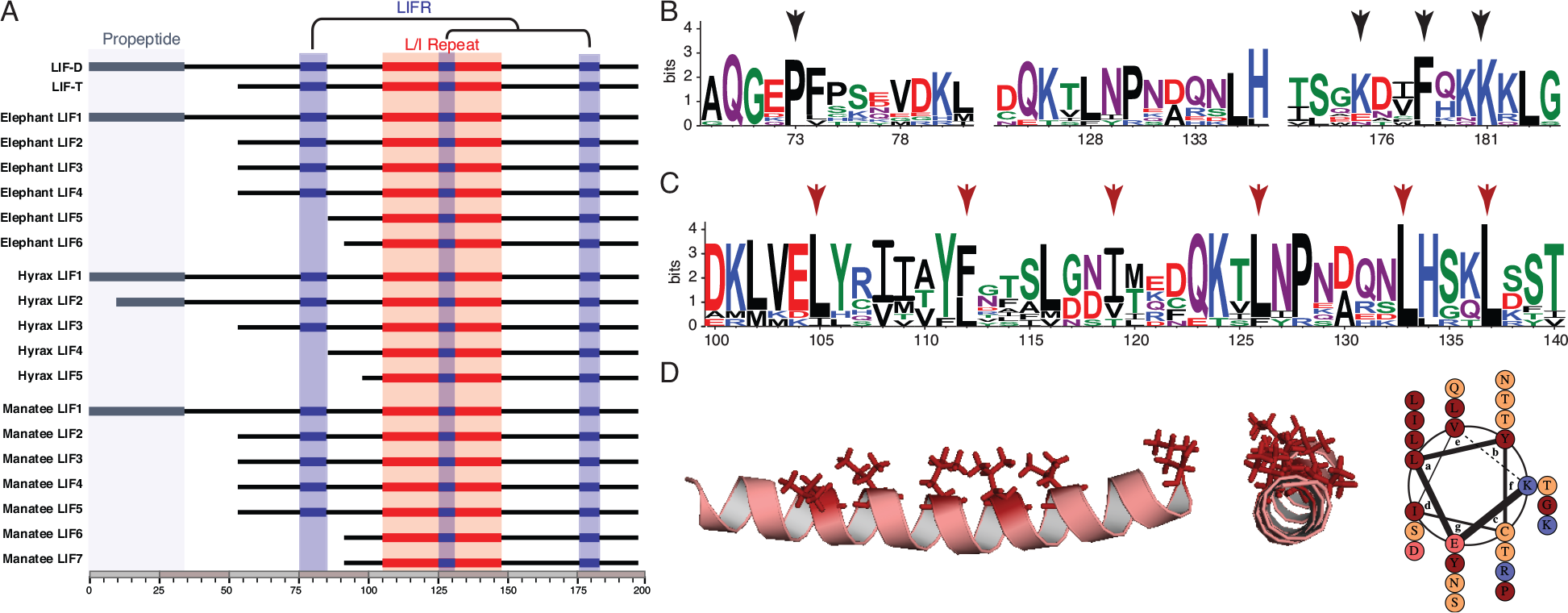
Structure of duplicate LIF genes with coding potential. (A) Domain structure of the LIF-D and LIF-T isoforms and of duplicate elephant, hyrax, and manatee LIF duplicates with coding potential. Locations of the propeptide, interactions sites with the LIF receptor (LIFR), and L/I repeat are shown. (B) Sequence logo showing conservation of LIF receptor (LIFR) interaction sites in duplicate LIF proteins. Residues in LIF that make physical contacts with LIFR are indicated with black arrows. Amino acids are colored according to physicochemical properties. Column height indicates overall conservation at that site (4, most conserved). (C) Sequence logo showing conservation of the leucine/isoleucine repeat region in duplicate LIF proteins. Leucine/isoleucine residues required for pro-apoptotic functions of LIF-T are indicated with red arrows. Amino acids are colored according to physicochemical properties. Column height indicates overall conservation at that site (4, most conserved). (D) Leucine/isoleucine residues in the African elephant LIF6 form an amphipathic alpha helix. Structural model of the LIF6 protein (left, center), and helical wheel representation of the LIF6 amphipathic alpha helix.

### Elephant *LIF6* is up-regulated by TP53 in response to DNA damage

If expansion of the *LIF* gene repertoire plays a role in the evolution of enhanced cancer resistance, then one or more of the *LIF* genes should be transcribed. To determine if duplicate *LIF* genes were transcribed, we assembled and quantified elephant *LIF* transcripts with HISAT2 (Kim et al., 2015) and StringTie (Pertea et al., 2015) using deep 100bp paired-end RNA-Seq data (>138 million reads) we previously generated from Asian elephant dermal fibroblasts (Sulak et al., 2016), as well as more shallow (~30 million reads) singe-end sequencing from Asian elephant peripheral blood mononuclear cells (PBMCs) (Reddy et al., 2015), African elephant dermal fibroblasts (Cortez et al., 2014) and placenta (Sulak et al., 2016). We identified transcripts corresponding to the LIF-D, LIF-M, and LIF-T isoforms of the canonical *LIF1* gene, and one transcript of a duplicate *LIF* gene (*LIF6*) in Asian elephant dermal fibroblasts (**Figure 4A**). The *LIF6* transcript initiates just downstream of canonical exon 2 and expression was extremely low (0.33 transcripts per million), as might be expected for a pro-apoptotic gene. No other RNA-Seq dataset identified duplicate LIF transcripts.

**Figure 4.**
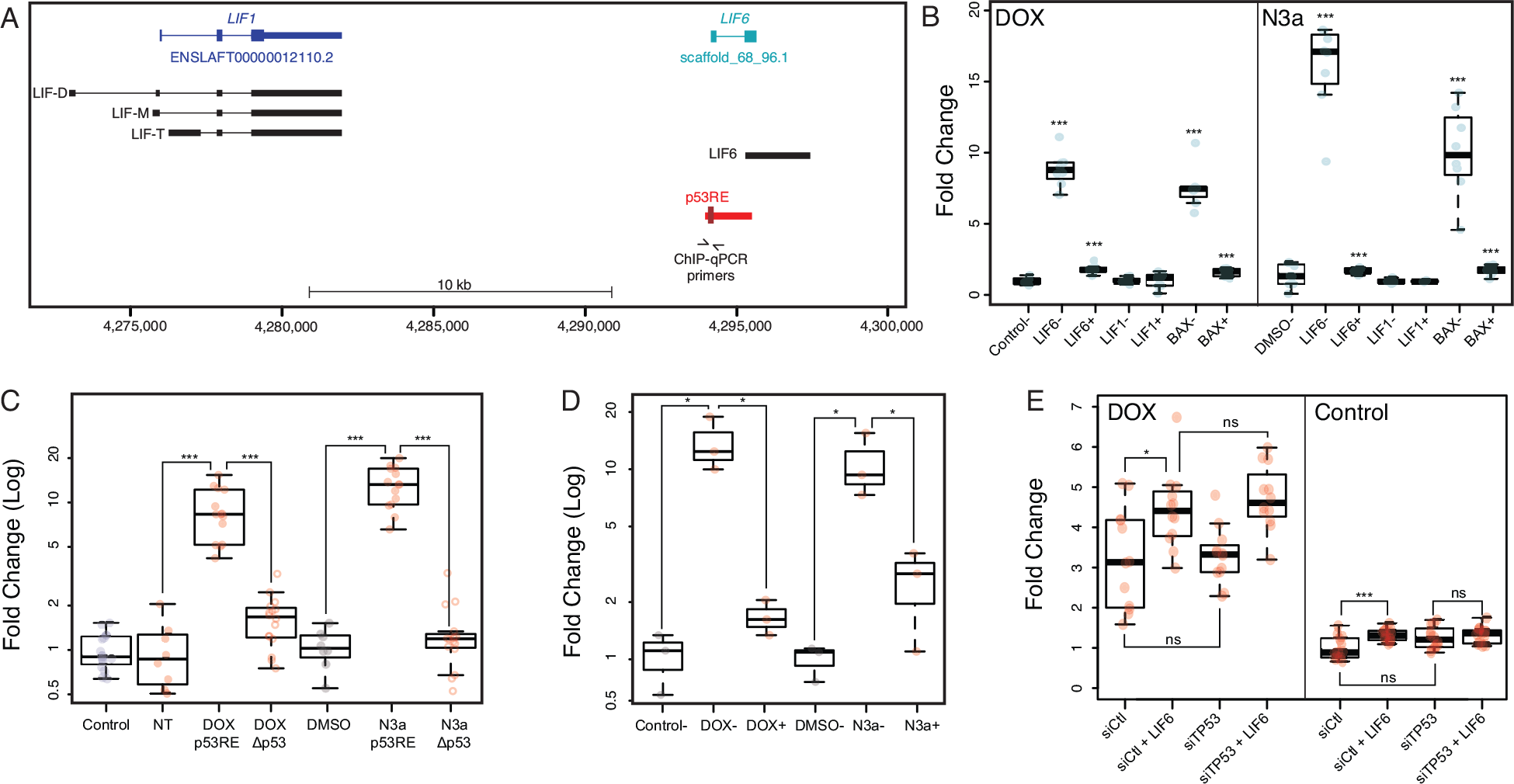
African elephant LIF6 is transcriptionally up-regualted by TP53 in response to DNA damage. (A) Structure of the African elephant *LIF/LIF6* locus (loxAfr3). The ENSEMBL LIF and geneID gene models are shown in blue and cyan. Transcripts assembled by StringTie (option ‘do not use GFF/GTF’) are shown in black. The region upstream of LIF6 used in transcription factor binding site prediction and luciferase assays is shown in red; the location of the putative p53 binding-site is shown in dark red. (B) Quantitative real-time PCR (qPCR) showing that *LIF6* is up-regulated in African elephant fibroblasts treated with doxorubicin (DOX) or nutlin-3a (N3a) and either a negative control siRNA (-) or an siRNA to knockdown TP53 expression (+); TP53 knockdown prevents *LIF6* up-regulation in response to DOX or N3a. Data shown as fold-change relative to control (water) or DMSO (a carrier for nutlin-3a). N=8, *** Wilcox test *P*<0.001. (C) Dual luciferase reporter assay indicates that the *LIF6* upstream region (p53RE) activates luciferase expression in African elephant fibroblasts treated in response to doxorubicin (DOX) or nutlin-3a treatment (N3a), and is significantly attenuated by deletion of the putative TP53 binding site (Ap53). Data shown as fold-change relative to controls (water for DOX, DMSO for N3a). NT, no DOX or nutlin-3a treatment. N=8, *** Wilcox test *P*<0.001. (D) ChIp-qPCR indicates that the putative TP53 binding site is bound by TP53 in response to in response to doxorubicin (DOX-) or nutlin-3a treatment (N3a-), and is significantly attenuated by siRNA mediated TP53 knockdown (DOX+ or N3a-). Data shown as fold-change relative to carrier controls (water or DMSO) and standardized to IgG control. N=3, * unequal variance T-test *P*<0.06. (E) Knockdown of TP53 inhibits DOX induced apoptosis in elephant African elephant fibroblasts. Fibroblasts were transiently transfected with either an negative control siRNA (siCtl) or three siRNAs targeting TP53, and either a empty vector control or a LIF6 expression vector. Apoptosis was assayed using an ApoTox-Glo 18 hours after treatment with DOX or control media. N=8, **** Wilcox test *P*<0.05, *** Wilcox test *P*<0.001.

Previous studies have shown that TP53 regulates basal and inducible transcription of *LIF* in response to DNA damage through a binding site located in *LIF* intron 1 (Baxter and Milner, 2010; Hu et al., 2007), suggesting that duplicate *LIF* genes may be regulated by TP53. Therefore we computationally predicted TP53 binding sites within a 3kb window around Atlantogenatan *LIF* genes and identified binding site motifs in the first intron of African elephant, hyrax, manatee, tenrec, and armadillo *LIF1* genes whereas the only duplicate *LIF* gene with a putative TP53 binding site was elephant *LIF6*; note that the putative TP53 binding sites around *LIF1* and *LIF6* are not homologous (**Figure S1**). Next we treated African elephant primary dermal fibroblasts with the DNA damaging agent doxorubicin (DOX) or the MDM2 antagonist nutlin-3a and quantified the transcription of canonical *LIF1*, duplicate *LIF* genes, and the TP53 target gene BAX by qRT-PCR. DOX treatment induced *LIF6* expression 8.18-fold (Wilcox test, *P*=1.54×10^−6^) and nutlin-3a induced *LIF6* expression 16.06-fold (Wilcox test, *P*=1.00×10^−4^), which was almost completely attenuated by siRNA mediated TP53 knockdown (**Figure S2** and **Figure 4B**). Treatment with DOX (Wilcox test, *P*=1.55×10^−4^) or nutlin-3a (Wilcox test, *P*=1.55×10^−4^) also up-regulated the TP53 target gene BAX (**Figure 4B**), which again was almost blocked by knockdown of TP53 (**Figure 4B**). In contrast neither treatment up-regulated *LIF1* (**Figure 4B**) and we observed no expression of the other duplicate *LIF* genes in African elephant fibroblasts or any *LIF* duplicate in hyrax fibroblasts treated with DOX or nutlin-3a. These data suggest that while *LIF6* encodes a transcribed gene in elephants, transcription of the other *LIF* duplicates is either induced by different signals or they are pseudogenes.

To test if the putative TP53 binding site upstream of elephant *LIF6* was a functional TP53 response element, we cloned the −1100bp to +30bp region of the African elephant *LIF6* gene into the pGL3-Basic[minP] luciferase reporter vector and tested its regulatory ability in dual luciferase reporter assays. We found that the African elephant *LIF6* upstream region had no effect on basal luciferase expression in transiently transfected African elephant fibroblasts (Wilcox test, *P*=0.53). In contrast, both DOX (Wilcox test, *P*=1.37×10^−8^) and nutlin-3a (Wilcox test, *P*=1.37×10^−8^) strongly increased luciferase expression (**Figure 4C**), which was almost completely abrogated by deletion of the putative TP53 binding-site in DOX (Wilcox test, *P*=1.37×10^−8^) and N3a (Wilcox test, *P*=1.37×10^−9^) treated cells (**Figure 4C**). Next we performed ChIP-qPCR to determine if the TP53 binding-site upstream of *LIF6* is bound by TP53 in African elephant fibroblasts treated with DOX or nutlin-3a using a rabbit polyclonal TP53 antibody (FL-393) that we previously demonstrated recognizes elephant TP53 (Sulak et al., 2016). DOX treatment increased TP53 binding 14.26-fold (unequal variance t-test, *P*=0.039) and nutlin-3a increased TP53 binding 10.75-fold (unequal variance t-test, *P*=0.058) relative to ChIP-qPCR with normal mouse IgG control antibody. This increased binding was almost completely attenuated by siRNA mediated TP53 knockdown (**Figure 4D**).

Finally, we transiently transfected elephant fibroblasts with either a negative control siRNA or siRNAs targeting *TP53* and a LIF6 expression vector and assayed cell viability, cytotoxicity, and apoptosis using an ApoTox-Glo assay 18 hours after treatment with DOX or control media. We found that LIF6 expression with negative control siRNAs augmented the induction of apoptosis by DOX (Wilcox test, *P*=0.033; **Figure 4E** and **Figure S3**). Knockdown of TP53 did not inhibit the induction of apoptosis (Wilcox test, *P*=0.033; **Figure 4E** and **Figure S3**), suggesting TP53 knockdown was insufficient to alter the induction of apoptosis; note that while siRNA mediated knockdown significantly reduced TP53 transcript levels (**Figure S2**), we were unable to validate knockdown of the TP53 protein because the FL-393 antibody that recognizes elephant TP53 is no longer available. Interestingly, however, LIF6 transfection induced apoptosis in elephant fibroblasts treated with control media and negative control siRNAs (Wilcox test, *P*=0.008), suggesting that LIF6 can induce apoptosis in the absence of DNA damage similar to LIF-T (**Figure 4E** and **Figure S3**). Thus, we conclude that elephant *LIF6* is transcriptionally up-regulated by TP53 in response to DNA damage and may have pro-apoptotic functions.

### Elephant LIF6 contributes to the augmented DNA-damage response in elephants

We have previously shown that elephant cells evolved to be extremely sensitive to genotoxic stress and induce apoptosis at lower levels of DNA damage than their closest living relatives, including the African Rock hyrax (*Procavia capensis capensis*), East African aardvark (*Orycteropus afer lademanni*), and Southern Three-banded armadillo (*Tolypeutes matacus*) (Sulak et al., 2016). To test the contribution of *LIF6* to this derived sensitivity, we designed a set of three siRNAs that specifically target *LIF6* and reduce *LIF6* transcript abundance ~88% (**Figure S2**). Next, we treated African elephant dermal fibroblasts with DOX or nutlin-3a and either *LIF6* targeting siRNAs or a control siRNA and assayed cell viability, cytotoxicity, and apoptosis using an ApoTox-Glo assay 24 hours after treatment. Both DOX (Wilcox test, *P*=3.33×10^−9^) and nutlin-3a (Wilcox test, *P*=3.33×10^−9^) reduced cell viability ~85%, which was attenuated 5-15% by *LIF6* knockdown in DOX (Wilcox test, *P*=1.33×10^−8^) or nutlin-3a (Wilcox test, *P*=3.33×10^−9^) treated cells (**Figure 5A**). While neither DOX nor nutlin-3a induced cytotoxicity (**Figure 5A**), both DOX (4.05-fold, Wilcox test, *P*=3.33×10^−9^) and nutlin-3a (2.64-fold, Wilcox test, *P*=3.33×10^−9^) induced apoptosis (**Figure 5A**).

**Figure 5.**
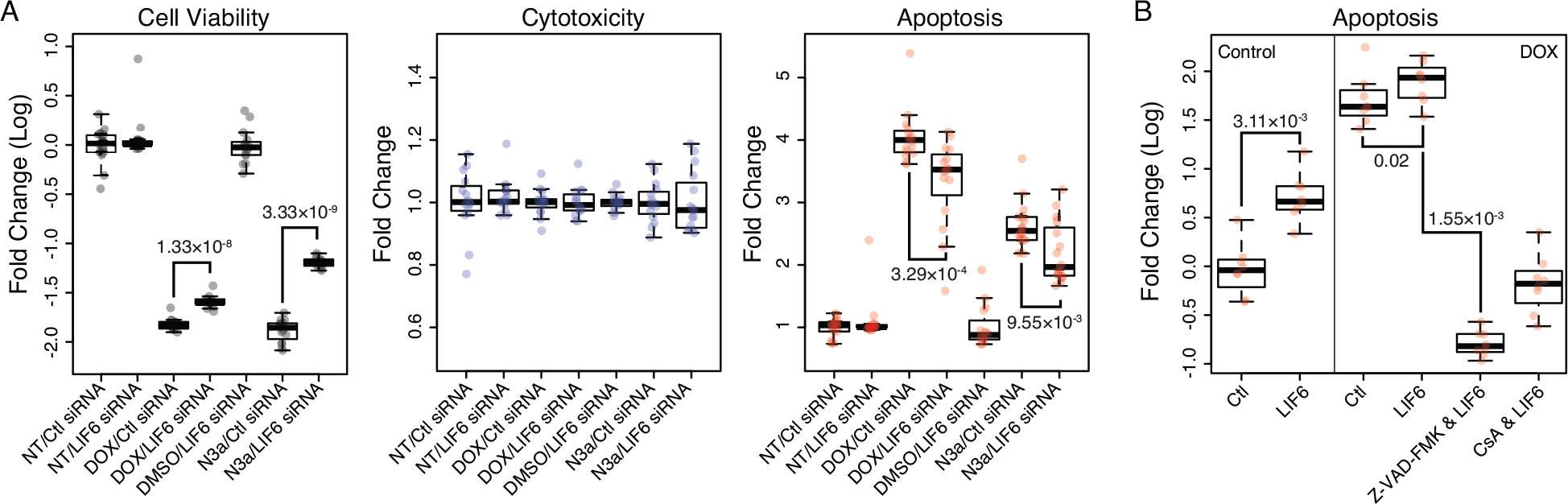
African elephant LIF6 contributes to the augmented DNA damage response in elephants. (A) African elephant fibroblasts were treated with either doxorubicin (DOX) or nutlin-3a (N3a), or an equimolar mixture of 3 siRNAs targeting LIF6 and doxorubicin (DOX/LIF6 siRNA) or nutlin-3a treatment (N3a/LIF6 siRNA). Cell viability, cytoxicity, and the induction of apoptosis was assayed using an ApoTox-Glo assay 24 hours after treatment. NT, no treatment. Ctl siRNA, negative control siRNA. DMSO, carrier for nutlin-3a. N=16, Wilcox test. (B)African elephant fibroblasts were transiently transfected with either an empty expression vector (Ctl) or a LIF6 encoding expression vector (LIF6), and treated with either DOX, the caspase inhibitor Z-VAD-FMK, or the cyclosporine A (CsA) which inhibits opening of the opening of the mitochondrial permeability transition pore. N=8, Wilcox test

To determine if LIF6 expression was sufficient to induce apoptosis, we transiently transfected a *LIF6* expression vector in to African elephant dermal fibroblasts and assayed cell viability, cytotoxicity, and apoptosis using the ApoTox-Glo assay 24 hours after transfection. We again found that LIF6 overexpression induced apoptosis in the absence of either DNA damage by DOX or TP53 activation by nutlin-3a treatment (Wilcox test, *P*=3.11×10^−4^), and augmented apoptosis induced with DOX (Wilcox test, *P*=0.02). Induction of apoptosis by LIF6 was almost completely blocked by co-treatment with the irreversible broad-spectrum caspase inhibitor Z-VAD-FMK (Wilcox test, *P*=1.55×10^−4^) but not cyclosporine A (Wilcox test, *P*=0.23), which inhibits opening of the opening of the mitochondrial permeability transition pore (**Figure 5B** and **Figure S4**). These data suggest that LIF6 contributes to the enhanced apoptotic response that evolved in the elephant lineage, likely through a mechanism that induces caspase-dependent apoptosis.

### Elephant LIF6 induces mitochondrial dysfunction and caspase-dependent apoptosis

To infer the mechanism(s) by which *LIF6* contributes to the induction of apoptosis, we first determined the sub-cellular localization of a LIF6-eGFP fusion protein in African elephant dermal fibroblasts. Unlike LIF-T, which has diffuse cytoplasmic and nuclear localization (Haines et al., 2000), LIF6-eGFP was located in discrete foci that co-localized with MitoTracker Red CM-H2XRos stained mitochondria (**Figure 6A**). Mitochondria are critical mediators of cell death, with distinct pathways and molecular effectors underlying death through either apoptosis (Karch et al., 2013; Tait and Green, 2010) or necrosis (Tait and Green, 2010; Vaseva et al., 2012). During apoptosis, for example, the Bcl-2 family members Bax/Bak form large pores in the outer mitochondrial membrane that allow cytochrome c to be released into the cytosol thereby activating the caspase cascade (Karch et al., 2013; Tait and Green, 2010). In contrast, during necrosis, Bax/Bak in the outer membrane interact with the cyclophilin D (CypD) and the inner membrane complex leading to the opening of the mitochondrial permeability transition pore (MPTP), swelling, and eventual rupture (Tait and Green, 2010; Vaseva et al., 2012).

**Figure 6.**
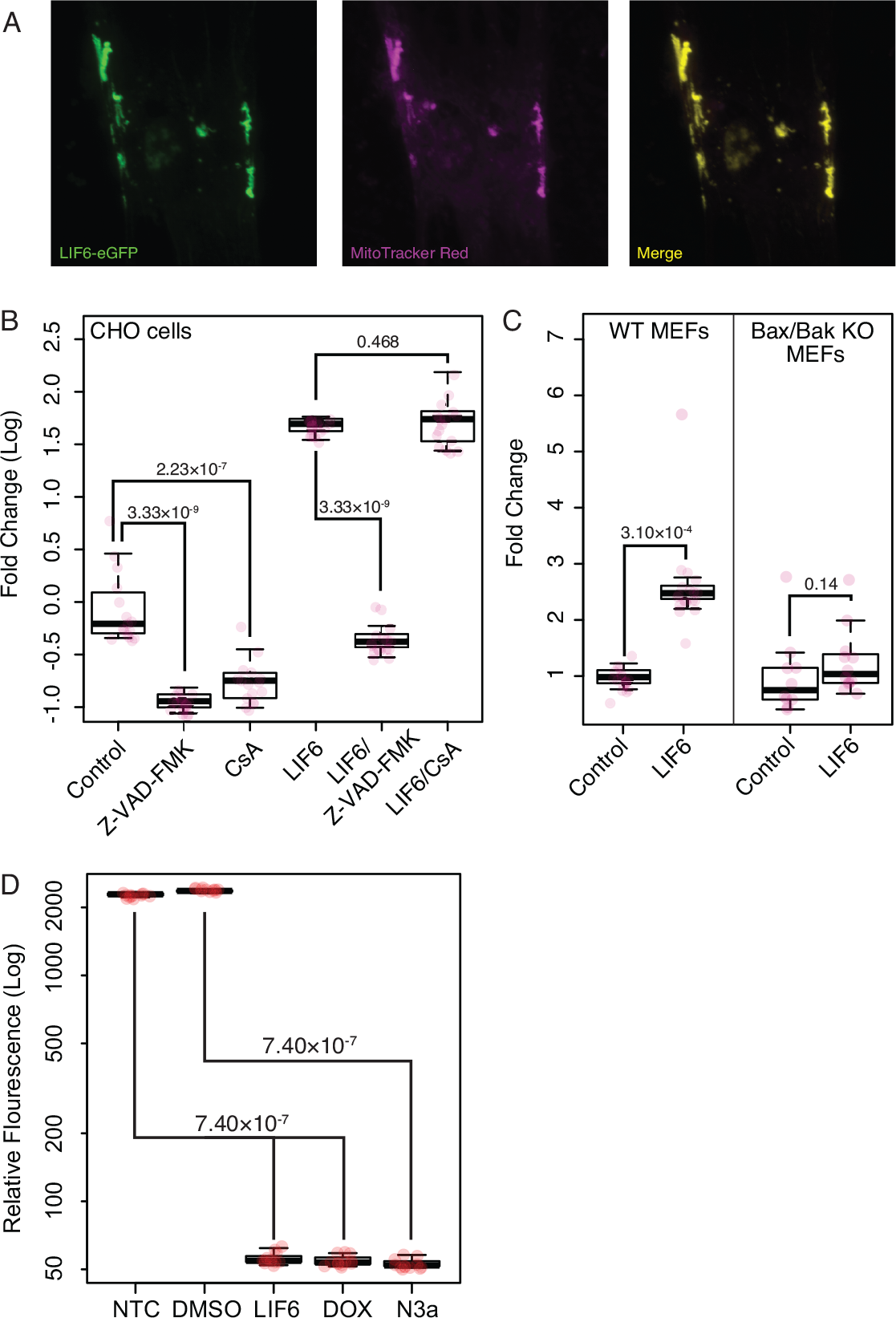
African elephant LIF6 is mitochondrial localized and induces caspase dependent apoptosis. (A) African elephant fibroblasts were transiently transfected with an expression vector encoding a eGFP tagged *LIF6* gene and mitochondria stained with MitoTracker Red CM-H2XRos. A single representative cell is shown. (B) Chinese hamster ovary (CHO) cells (which do not express LIFR) were transiently transfected with an expression vector encoding the African elephant *LIF6* gene and assayed for the induction of apoptosis with an ApoTox-Glo assay 24 hours after transfection. Induction of apoptosis by LIF6 was inhibited by co-treatment with the irreversible broad-spectrum caspase inhibitor Z-VAD-FMK but not cyclosporine-A (CsA). Treatment of CHO cells with Z-VAD-FMK or CsA alone reduced apoptosis. N=16, Wilcox test. (C) Overexpression of LIF6 in Bax/Bak double knockout mouse embryonic fibroblasts does not induce apoptosis, not augmented nutlin-3a induced apoptosis. N=8, Wilcox test. (D) Overexpression of LIF6 in CHO cells induces loss of mitochondrial membrane potential 48 hours after transfection. N=8, Wilcox test.

To test if LIF6 induced apoptosis was specific to elephant cells and independent of LIF receptor (LIFR) mediated signaling, we transiently transfected Chinese hamster (*Cricetulus griseus*) ovary (CHO) cells, which do not express *LIFR* (Orellana et al., 2018), with the LIF6 expression vector and assayed the induction of apoptosis with the ApoTox-Glo assay. Overexpression of LIF6 induced apoptosis 5.38-fold (Wilcox test, *P*=3.33×10^−9^) 24 hours after transfection, consistent with a pro-apoptotic function independent of LIFR (**Figure 6B**). Induction of apoptosis by LIF6, however, was almost completely blocked by co-treatment with Z-VAD-FMK (**Figure 6B**) but not cyclosporine A (CsA) (**Figure 6B**). To test if LIF6 induced apoptosis is dependent upon Bax and Bak, we overexpressed LIF6 in Bax/Bak knockout mouse embryonic fibroblasts (MEFs) but did not observe an induction of apoptosis (Wilcox test, *P*=0.14; **Figure 6C** and **Figure S5**). In contrast LIF6 overexpression induced apoptosis in wild-type MEFs (Wilcox test, *P*=0. 3.10×10^−4^; **Figure 6C** and **Figure S5**). During apoptosis, collapse of the mitochondrial membrane potential (MMP) coincides with the opening of the mitochondrial transition pores, leading to the release of proapoptotic factors into the cytosol. Consistent with this mechanism, we found that LIF6 overexpression, treatment with DOX, or with nutlin-3a induced loss of MMP in CHO cells 48 hours after transfection (Wilcox test, *P*=7.40×10^−7^; **Figure 6D**). Thus *LIF6* is sufficient to induce mitochondrial dysfunction and apoptosis mediated through Bax/Bak and independent of MPTP opening.

### Elephant *LIF6* is a refunctionalized pseudogene

We reasoned that most duplicate *LIF* genes are (likely) pseudogenes because elephant *LIF6* is deeply nested within the duplicate *LIF* clade, is the only expressed duplicate, and is the only duplicate with a TP53 response element, suggesting elephant *LIF6* re-evolved into a functional gene from a pseudogene ancestor. To test this hypothesis and reconstruct the evolutionary history of the *LIF6* gene in the Proboscideans with greater phylogenetic resolution, we annotated the *LIF6* locus in the genomes of Elephantids including the African Savannah elephant (*Loxodonta africana*), African Forest elephant (*Loxodonta cyclotis*), Asian elephant (*Elephas maximus*), woolly mammoth (*Mammuthus primigenius*), Columbian mammoth (*Mammuthus columbi*), and straight-tusked elephant (*Palaeoloxodon antiquus*), as well as the American Mastodon (*Mammut americanum*), an extinct Mammutid. We found that the genomes of each extinct Proboscidean contained a *LIF6* gene with coding potential similar to the African and Asian elephant *LIF6* genes as well as the TP53 binding-site, indicating that *LIF6* evolved to be a TP53 target gene in the stem-lineage of Proboscideans.

While functional genes evolve under selective constraints that reduce their *d_N_*/*d_S_* (ω) ratio to below one, pseudogenes are generally free of such constraints and experience a relaxation in the intensity of purifying selection and an elevation in their *d_N_*/*d_S_* ratio. Therefore, we used a random effects branch-site model (RELAX) to test for relaxed selection on duplicate *LIF* genes compared to canonical *LIF* genes. The RELAX method fits a codon model with three ω rate classes to the phylogeny (null model), then tests for relaxed/intensified selection along lineages by incorporating a selection intensity parameter (K) to the inferred ω values; relaxed selection (both positive and negative) intensity is inferred when K<1 and increased selection intensity is inferred when K>1. As expected for pseudogenes, *LIF* duplicates (other than Proboscidean *LIF6* genes) had significant evidence for a relaxation in the intensity of selection (K=0.36, LRT=42.19, *P*=8.26×10^−11^) as did the Proboscidean *LIF6* stem-lineage (K=0.00, LRT=3.84, *P*=0.05). In contrast, Proboscidean *LIF6* genes had significant evidence for selection intensification (K=50, LRT=4.46, *P*=0.03). We also found that the branch-site unrestricted statistical test for episodic diversification (BUSTED), which can detect gene-wide (not site-specific) positive selection on at least one site and on at least one branch, inferred a class of strongly constrained sites in (ω=0.00, 23.74%), a class of moderately constrained sites (ω=0.64, 75.85%), and a few sites that may have experienced positive selection in Proboscidean *LIF6* genes (ω=10000.00, 0.41%; LRT=48.81, *P*≤0.001). These data are consistent with the reacquisition of constraints after refunctionalization.

Finally we inferred a Bayesian time-calibrated phylogeny of Atlantogenatan *LIF* genes, including *LIF6* from African and Asian elephant, woolly and Columbian mammoth, straight-tusked elephant, and American Mastodon, to place upper and lower bounds on when the Proboscidean *LIF6* gene may have refunctionalized (**Figure 7A**). We found that estimated divergence date of the Proboscideans *LIF6* lineage was ~59 MYA (95% HPD: 61-57 MYA) whereas the divergence of Proboscideans was ~26 MYA (95% HPD: 23.28 MYA). These data indicate that the Proboscidean *LIF6* gene refunctionalized during the evolutionary origin of large body sizes in this lineage, although precisely when within this time interval is unclear (**Figure 7B**). Thus LIF6 was reanimated sometime before the demands of maintaining a larger body existed in the Proboscidean lineage, suggesting LIF6 is permissive for the origin of large bodies but is not sufficient.

**Figure 7.**
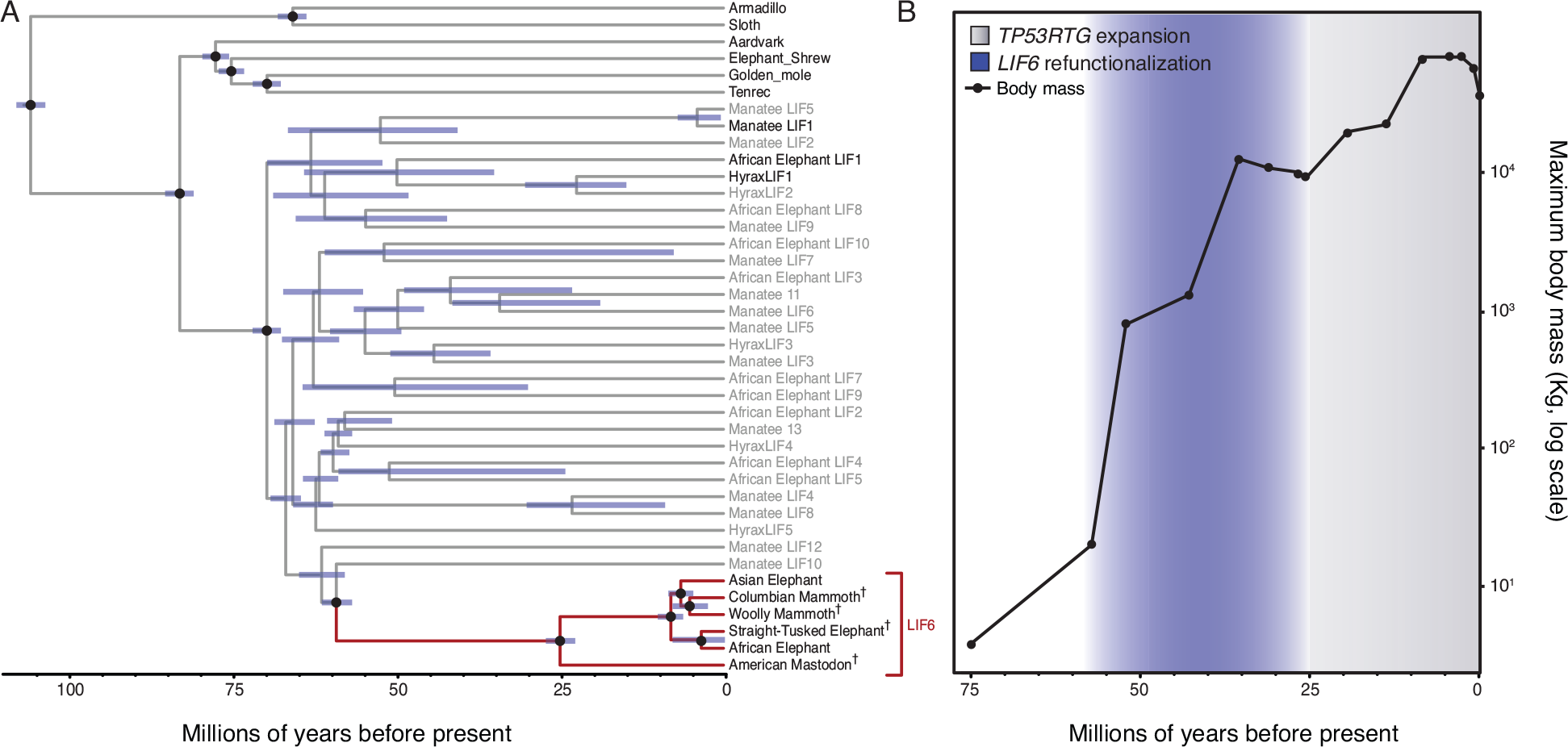
LIF6 is a re-functionalized pseudogene. (A) Time calibrated Bayesian phylogeny of Atlantogenatan LIF genes. The Proboscidean LIF6 clade is highlighted in red, canonical LIF genes in black, LIF duplicates in grey. The 95% highest posterior density (HPD) of estimated divergence dates are shown as blue bars. Nodes used to calibrate divergence dates are shown with black circles. (B) Proboscidean LIF6 re-functionalized during the evolution of large body sizes in the Proboscidean lineage.

## Discussion

Identifying the molecular and genetic mechanisms that underlie the evolution of enhanced cancer resistance in large and long-lived species is important for understanding how animals resolved Peto’s paradox and may provide a better understanding of cancer biology, but has been hampered by a lack of genomes for species that differ dramatically in body size yet are relatively recently diverged. Fortunately sequenced genomes are available for several living and extinct elephant species including the African Savannah elephant, African Forest elephant, Asian elephant, woolly and Columbian mammoths, straight-tusked elephant, the American Mastodon, an extinct Mammutid that diverged from the elephant lineage ~25 million years ago, as well as manatee and rock hyrax which are the closest living relatives of Proboscideans. This allowed us to use comparative genomic methods to identify genetic changes that occurred during the evolution of large bodies in the Proboscidean lineage, such as gene duplication and loss events.

A comprehensive analyses of genetic changes associated with the resolution of Peto’s paradox in the elephant lineage has yet to be performed, but candidate gene studies have identified functional duplicates of the master tumor suppressor TP53 as well as putative duplicates of other tumor suppressor genes (Abegglen et al., 2015; Caulin et al., 2015; Sulak et al., 2016). Caulin et al, for example, characterized the copy number of 830 tumor-suppressor genes (Higgins et al., 2007) across 36 mammals and identified 382 putative duplicates, including five copies of *LIF* in African elephants, seven in hyrax, and three in tenrec. Here we show that an incomplete duplication of the *LIF* gene in the Paenungulate stem-lineage generated a duplicate missing the proximal promoter and exon 1, generating a gene with similar structure to the LIF-T isoform (Haines et al., 1999), which functions as an intra-cellular pro-apoptotic protein independently from the LIFR-mediated signaling. Additional duplications of this original duplicate increased *LIF* copy number in Paenungulates, however, most *LIF* duplicates lack regulatory elements, are not expressed in elephant or hyrax fibroblasts (manatee cells or tissues are unavailable), and, with the exception of elephant *LIF6*, are likely pseudogenes.

While we are unable to do the kinds of reverse and forward genetic experiments that traditionally establish causal associations between genotypes and phenotypes, we were able to use primary African elephant and hyrax dermal fibroblasts to functionally characterize *LIF* duplicates. We found, for example, that the elephant *LIF6* gene is transcribed at very low levels under basal conditions, but is up-regulated by TP53 in response to DNA damage. One of the constraints on the refunctionalization of pseudogenes is that they must evolve new cis-regulatory elements to direct their expression, but random DNA sequences can evolve into promoters with only a few substitutions suggesting de novo origination of regulatory elements may be common (Yona et al., 2017). There should be strong selection against the origin of constitutively active enhancers/promoters for pro-apoptotic pseudogenes, however, because their expression will be toxic. These results imply refunctionalizing *LIF* pseudogenes may impose a potential evolutionary cost. One of the ways to avoid that cost is through the gain of inducible regulatory elements that appropriately respond to specific stimuli, such as a TP53 signaling. Indeed our phylogenetic analysis indicates that a TP53 response element up-stream of *LIF6* evolved before the divergence of mastodons and the modern elephant lineage, suggesting that *LIF6* refunctionalized in the stem-lineage of Proboscideans coincident with the origin of large body sizes and thus may have been permissive for the large bodies.

The precise mechanisms by which mitochondrial dysfunction leads to apoptosis are uncertain, however, during early stages of apoptosis the pro-death Bcl-2 family members Bax and Bak hetero- and homo-oligomerize within the mitochondrial outer membrane leading to permeabilization (MOMP) and the release of pro-apoptotic protein such as cytochrome c (Karch et al., 2013; 2015). In contrast, during necrosis the collapse of the MMP and the opening of the mitochondrial permeability transition pore (MPTP) leads to mitochondrial swelling, rupture, and cell death (Ly et al., 2003). Our observations that cyclosporine A (CsA) did not inhibit LIF6 induced apoptosis, and that LIF6 overexpression did not induce apoptosis in Bax/Bak null MEFs suggests that LIF6 functions in a manner analogous to the pro-apoptotic Bcl-2 family members by inducing the opening of the outer mitochondrial membrane pore. Furthermore our observation that LIF6 overexpression incudes apoptosis in elephant dermal fibroblasts, Chinese hamster ovary cells, and mouse embryonic fibroblasts indicates the LIF6 mechanism of action is neither of cell-type nor species specific. The molecular mechanisms by which LIF6 induces apoptosis, however, are unclear and the focus of continued studies.

## Experimental Procedures

### Identification of *LIF* genes in Mammalian genomes

We used BLAT to search for *LIF* genes in 53 Sarcopterygian genomes using the human LIF protein sequences as an initial query. After identifying the canonical *LIF* gene from each species, we used the nucleotide sequences corresponding to this *LIF* CDS as the query sequence for additional BLAT searches within that species genome. To further confirm the orthology of each *LIF* gene we used a reciprocal best BLAT approach, sequentially using the putative CDS of each *LIF* gene as a query against the human genome; in each case the query gene was identified as *LIF*. Finally we used the putative amino acid sequence of the LIF protein as a query sequence in a BLAT search.

We thus used BLAT to characterize the *LIF* copy number in Human (*Homo sapiens*; GRCh37/hg19), Chimp (*Pan troglodytes*; CSAC 2.1.4/panTro4), Gorilla (*Gorilla gorilla gorilla*; gorGor3.1/gorGor3), Orangutan (*Pongo pygmaeus abelii*; WUGSC 2.0.2/ponAbe2), Gibbon (*Nomascus leucogenys*; GGSC Nleu3.0/nomLeu3), Rhesus (*Macaca mulatta*; BGI CR_1.0/rheMac3), Baboon (*Papio hamadryas*; Baylor Pham_1.0/papHam1), Marmoset (*Callithrix jacchus*; WUGSC 3.2/calJac3), Squirrel monkey (*Saimiri boliviensis*; Broad/saiBol1), Tarsier (*Tarsius syrichta*; Tarsius_syrichta2.0.1/tarSyr2), Bushbaby (*Otolemur garnettii*; Broad/otoGar3), Mouse lemur (*Microcebus murinus*; Broad/micMur1), Chinese tree shrew (*Tupaia chinensis*; TupChi_1.0/tupChi1), Squirrel (*Spermophilus tridecemlineatus*; Broad/speTri2), Mouse (*Mus musculus*; GRCm38/mm10), Rat (*Rattus norvegicus*; RGSC 5.0/rn5), Naked mole-rat (*Heterocephalus glaber*; Broad HetGla_female_1.0/hetGla2), Guinea pig (*Cavia porcellus*; Broad/cavPor3), Rabbit (*Oryctolagus cuniculus*; Broad/oryCun2), Pika (*Ochotona princeps*; OchPri3.0/ochPri3), Kangaroo rat (*Dipodomys ordii*; Broad/dipOrd1), Chinese hamster (*Cricetulus griseus*; C_griseus_v1.0/criGri1), Pig (*Sus scrofa*; SGSC Sscrofa10.2/susScr3), Alpaca (*Vicugna pacos*; Vicugna_pacos-2.0.1/vicPac2), Dolphin (*Tursiops truncatus*; Baylor Ttru_1.4/turTru2), Cow (*Bos taurus*; Baylor Btau_4.6.1/bosTau7), Sheep (*Ovis aries*; ISGC Oar_v3.1/oviAri3), Horse (*Equus caballus*; Broad/equCab2), White rhinoceros (*Ceratotherium simum*; CerSimSim1.0/cerSim1), Cat (*Felis catus*; ICGSC Felis_catus 6.2/felCat5), Dog (*Canis lupus familiaris*; Broad CanFam3.1/canFam3), Ferret (*Mustela putorius furo*; MusPutFur1.0/musFur1), Panda (*Ailuropoda melanoleuca*; BGI-Shenzhen 1.0/ailMel1), Megabat (*Pteropus vampyrus*; Broad/pteVam1), Microbat (*Myotis lucifugus*; Broad Institute Myoluc2.0/myoLuc2), Hedgehog (*Erinaceus europaeus*; EriEur2.0/eriEur2), Shrew (*Sorex araneus*; Broad/sorAra2), Minke whale (*Balaenoptera acutorostrata scammoni*; balAcu1), Bowhead Whale (*Balaena mysticetus*; v1.0), Rock hyrax (*Procavia capensis*; Broad/proCap1), Sloth (*Choloepus hoffmanni*; Broad/choHof1), Elephant (*Loxodonta africana*; Broad/loxAfr3), Cape elephant shrew (*Elephantulus edwardii*; EleEdw1.0/eleEdw1), Manatee (*Trichechus manatus latirostris*; Broad v1.0/triMan1), Tenrec (*Echinops telfairi*; Broad/echTel2), Aardvark (*Orycteropus afer afer*; OryAfe1.0/oryAfe1), Armadillo (*Dasypus novemcinctus*; Baylor/dasNov3), Opossum (*Monodelphis domestica*; Broad/monDom5), Tasmanian devil (*Sarcophilus harrisii*; WTSI Devil_ref v7.0/sarHar1), Wallaby (*Macropus eugenii*; TWGS Meug_1.1/macEug2), and Platypus (*Ornithorhynchus anatinus*; WUGSC 5.0.1/ornAna1).

### Phylogenetic analyses and gene tree reconciliation of Paenungulate *LIF* genes

The phylogeny of *LIF* genes was estimated using an alignment of the *LIF* loci from the African elephant, hyrax, manatee, tenrec, and armadillo genomes and BEAST (v1.8.3) (Rohland et al., 2010). We used the HKY85 substitution, which was chosen as the best model using HyPhy, empirical nucleotide frequencies (+F), a proportion of invariable sites estimated from the data (+I), four gamma distributed rate categories (+G), an uncorrelated random local clock to model substitution rate variation across lineages, a Yule speciation tree prior, uniform priors for the GTR substitution parameters, gamma shape parameter, proportion of invariant sites parameter, and nucleotide frequency parameter. We used an Unweighted Pair Group Arithmetic Mean (UPGMA) starting tree. The analysis was run for 10 million generations and sampled every 1000 generations with a burn-in of 1000 sampled trees; convergence was assessed using Tracer, which indicated convergence was reached rapidly (within 100,000 generations). We used Notung v2.6 (Chen et al., 2000) to reconcile the gene and species trees.

### Gene expression data (Analyses of RNA-Seq data and RT-PCR)

To determine if duplicate *LIF* genes were basally transcribed, we assembled and quantified elephant *LIF* transcripts with HISAT2 (Kim et al., 2015) and StringTie (Pertea et al., 2015) using deep 100bp paired-end RNA-Seq data (>138 million reads) we previously generated from Asian elephant dermal fibroblasts (Sulak et al., 2016), as well as more shallow (~30 million reads) singe-end sequencing from African elephant dermal fibroblasts (Cortez et al., 2014) and placenta (Sulak et al., 2016), and Asian elephant peripheral blood mononuclear cells (PBMCs) (Reddy et al., 2015). HISAT2 and StringTie were run on the Galaxy web-based platform (https://usegalaxy.org) (Afgan et al., 2016) using default settings, and without a guide GTF/GFF file.

We determined if *LIF* transcription was induced by DNA damage and p53 activation in African elephant Primary fibroblasts (San Diego Frozen Zoo) using RT-PCR and primers designed to amplify elephant duplicate *LIF* genes, including LIF1-F: 5’-GCACAGAGAAGGACAAGCTG-3’, LIF1-R: 5’-CACGTGGTACTTGTTGCACA-3’, LIF6-F: 5’-CAGCTAGACTTCGTGGCAAC-3’, LIF6-R: 5’-AGCTCAGTGATGACCTGCTT-3’, LIF3-R: 5’-TCTTTGGCTGAGGTGTAGGG-3’, LIF4-F: 5’-GGCACGGAAAAGGACAAGTT-3’, LIF4-R: 5’-GCCGTGCGTACTTTATCAGG-3’, LIF5-F: 5’-CTCCACAGCAAGCTCAAGTC-3’, LIF5-R: 5’-GGGGATGAGCTGTGTGTACT-3’. We also used primers to elephant BAX to determine if it was up-regulated by TP53: BAX-F: 5’-CATCCAGGATCGAGCAAAGC-3’, BAX-R: 5’-CCACAGCTGCAATCATCCTC-3’. African elephant Primary fibroblasts were grown to 80% confluency in T-75 culture flasks at 37°C/5% CO_2_ in a culture medium consisting of FGM/EMEM (1:1) supplemented with insulin, FGF, 6% FBS and Gentamicin/Amphotericin B (FGM-2, singlequots, Clonetics/Lonza). At 80% confluency, cells were harvested and seeded into 6-well culture plates at ~10,000 cells/well. Once cells recovered to 80% confluency they were treated with either vehicle control, 50um Doxorubicin, or 50um Nutlin-3a.

Total RNA was extracted using the RNAeasy Plus Mini kit (Qiagen), then DNase treated (Turbo DNA-*free* kit, Ambion) and reverse-transcribed using an olgio-dT primer for cDNA synthesis (Maxima H Minus First Strand cDNA Synthesis kit, Thermo Scientific). Control RT reactions were otherwise processed identically, except for the omission of reverse transcriptase from the reaction mixture. RT products were PCR-amplified for 45 cycles of 94°/20 seconds, 56°/30 seconds, 72°/30 seconds using a BioRad CFX96 Real Time qPCR detection system and SYBR Green master mix (QuantiTect, Qiagen). PCR products were electrophoresed on 3% agarose gels for 1 hour at 100 volts, stained with SYBR safe, and imaged in a digital gel box (ChemiDoc MP, BioRad) to visualize relative amplicon sizes.

### Statistical methods

We used a Wilcox or T-test test for all statistical comparisons, with at least four biological replicates. The specific statistical test used and number replicates for each experiment are indicated in figure legends.

### Luciferase assay and cell culture

We used the JASPAR database of transcription factor binding site (TFBS) motifs (Mathelier et al., 2015) to computationally predict putative TFBSs within a 3kb window around Atlantogenatan *LIF* genes and identified matches for the TP53 motif (MA0106.3), including a match (sequence: CACATGTCCTGGCAACCT, score: 8.22, relative score: 0.82) ~1kb upstream of the African elephant *LIF6* start codon. To test if the putative p53 binding site upstream of elephant *LIF6* was a functional p53 response element, we synthesized (GeneScript) and cloned the −1100bp to +30bp region of the African elephant *LIF6* gene (loxAfr3_dna range=scaffold_68:4294134-4295330 strand=+ repeatMasking=none) and a mutant lacking the CACATGTCCTGGCAACCT sequence into the pGL3-Basic[minP] luciferase reporter vector.

African elephant Primary fibroblasts (San Diego Frozen Zoo) were grown to 80% confluency in T-75 culture flasks at 37°C/5% CO_2_ in a culture medium consisting of FGM/EMEM (1:1) supplemented with insulin, FGF, 6% FBS and Gentamicin/Amphotericin B (FGM-2, singlequots, Clonetics/Lonza). At 80% confluency, 10^4^ cells were harvested and seeded into 96-well white culture plates. 24 hours later cells were transfected using Lipofectamine LTX and either 100g of the pGL3-Basic[minP], pGL3-Basic[minP] −1100bp to +30bp, pGL3-Basic[minP] - 1100bp-+30bp Δp53TFBS luciferase reporter vectors and 1ng of the pGL4.74[*hRluc*/TK] Renilla control reporter vector according the standard protocol with 0.5 ul/well of Lipofectamine LTX Reagent and 0.1ul/well of PLUS Reagent. 24 hours after transfection cells were treated with either vehicle control, 50um Doxorubicin, or 50um Nutlin-3a. Luciferase expression was assayed 48 hours after drug treatment, using the Dual-Luciferase Reporter Assay System (Promega) in a GloMax-Multi+ Reader (Promega). For all experiments luciferase expression was standardized to Renilla expression to control for differences transfection efficiency across samples; Luc./Renilla data is standardized to (Luc./Renilla) expression in untreated control cells. Each luciferase experiment was replicated three independent times, with 8-16 biological replicates per treatment and control group.

### ChIP-qPCR and cell culture

African elephant Primary fibroblasts were grown to 80% confluency in T-75 culture flasks at 37°C/5% CO_2_ in a culture medium consisting of FGM/EMEM (1:1) supplemented with insulin, FGF, 6% FBS and Gentamicin/Amphotericin B (FGM-2, singlequots, Clonetics/Lonza). 10^4^ cells were seeded into each well of 6-well plate and grown to ~80% confluency. Cells were then treated with either a negative control siRNA or equimolar amounts of a combination of three siRNAs that specifically target the canonical TP53 transcript using Lipofectamine LTX according to the suggested standard protocol. The next day, cells were treated with either water, DMSO, 50uM Doxorubicin, or 50uM Nutlin-3a in three biological replicates for each condition. After 18 hrs of incubation with each drug, wells were washed three times with ice cold PBS and PBS replaced with fresh media, and chromatin cross linked with 1% fresh formaldehyde for 10 minutes. We used The MAGnify Chromatin Immunoprecipitation System (ThermoFischer #492024) to perform chromatin immunoprecipitation according to the suggested protocol. However rather than shearing chromatin by sonication, we used the ChIP-It Express Enzymatic Shearing Kit (Active Motif # 53009) according to the suggested protocol. Specific modifications to the MAGnify Chromatin Immunoprecipitation System included using 3ug of the polyclonal TP53 antibody (FL-393, lot #DO215, Santa Cruz Biotechnology).

We used qPCR to assay for enrichment of TP53 binding from the ChIP-Seq using the forward primer 5’-TGGTTTCCAGGAGTCTTGCT-3’ and the reverse primer 5’-CATCCCCTCCTTCCTCTGTC-3’. 100ng of ChIP DNA was used per PCR reaction, which was amplified for 45 cycles of 94°/20 s, 56°/30 s, 72°/30 s using a BioRad CFX96 Real Time qPCR detection system and SYBR Green master mix (QuantiTect, Qiagen). Data are shown as fold increase in TP53 ChIP signal relative to the background rabbit IgG ChIP signal and standardized to the control water for DOX or DMSO for nutlin-3a treatments.

### ApoTox-Glo Viability/Cytotoxicity/Apoptosis experiments

T75 culture flasks were seeded with 200,000 African Elephant primary fibroblasts, and grown to 80% confluency at 37°C/5% CO2 in a culture medium consisting of FGM/EMEM (1:1) supplemented with insulin, FGF, 6% FBS and Gentamicin/Amphotericin B (FGM-2, singlequots, Clonetics/Lonza). 5000 cells were seeded into each well of two opaque-bottomed 96-well plates. In each plate, half of the columns in the plate were transfected with pcDNA3.1/LIF6/eGFP (GenScript) using Lipofectamine LTX (Thermo Scientific 15338100); the other half were mock transfected with the same protocol without any DNA. In the plate designated for the 18hr timepoint, each column was treated with either: 50uM (-)-Nutlin-3 (Cayman 18585); 20uM Z-VAD-FMK (Cayman 14463); 2uM Cyclosporin A (Cayman 12088); 50uM Doxorubicin (Fisher BP251610); DMSO (Fisher BP231100); or DPBS (Gibco 14190136). For the 24hr timepoint, the same schema for treatment was used, but with half-doses. Each treatment contained eight biological replicates for each condition. After 18 hrs of incubation with each drug, cell viability, cytotoxicity, and Caspase-3/7 activity were measured using the ApoTox-Glo Triplex Assay (Promega) in a GloMax-Multi+ Reader (Promega). Z-VAD-FMK readings were normalized to the PBS-treated, mock-transfected cells; all others were normalized to the DMSO-treated, mock-transfected cells.

T75 culture flasks were seeded with 250,000 wild-type (ATCC CRL-2907) and Bak/Bax double knockout (ATCC CRL-2913) mouse embryonic fibroblasts (MEFs), or Chinese hamster ovary cells (CHO-K1, Thermo R75807) and allowed to grow to 80% confluency at 37°C/5% CO2 in a culture medium consisting of high-glucose DMEM (Gibco) supplemented with GlutaMax (Gibco), Sodium pyruvate (Gibco), 10% FBS (Gibco), and Penicillin-Streptomycin (Gibco). 3000 cells were seeded into each well of an opaque, bottomed 96-well plate. Half of the columns in the plate were transfected with pcDNA3.1/LIF6/eGFP (GenScript) using Lipofectamine LTX (ThermoFisher Scientific 15338100); the other half were mock transfected with the same protocol without any DNA. 6 hours post-transfection, the transfection reagents and media from each well was replaced: for the 24-hour timepoint, drug-supplemented media was placed within the wells; for the 48-hour timepoint, untreated media was placed in the wells, and then replaced with treatment media 24-hours later. Each column was treated with either: 50uM (-)-Nutlin-3 (Cayman 18585); 20uM Z-VAD-FMK (Cayman 14463); 2uM Cyclosporin A (Cayman 12088); 50uM Doxorubicin (Fisher BP251610); DMSO (Fisher BP231100); or DPBS (Gibco 14190136). Each treatment contained eight biological replicates for each condition. After 18 hrs of incubation with each drug, cell viability, cytotoxicity, and Caspase-3/7 activity were measured using the ApoTox-Glo Triplex Assay (Promega) in a GloMax-Multi+ Reader (Promega). Z-VAD-FMK readings were normalized to the PBS-treated, mock-transfected cells; all others were normalized to the DMSO-treated, mock-transfected cells.

For knockdown experiments T75 culture flasks were seeded with 200,000 African Elephant primary fibroblasts, and grown to 80% confluency at 37°C/5% CO2 in a culture medium consisting of FGM/EMEM (1:1) supplemented with insulin, FGF, 6% FBS and Gentamicin/Amphotericin B (FGM-2, singlequots, Clonetics/Lonza). 5000 cells were seeded into each well of two opaque-bottomed 96-well plates. In each plate, pairs of rows were transfected with either Silencer™ Select Negative Control No. 1 siRNA (Thermo 4390843), P53 siRNA (Dharmacon) (Sulak et al. 2016), and either with or without pcDNA3.1/LIF6/eGFP (GenScript) using Lipofectamine LTX (Thermo Scientific 15338100). In the plate designated for the 18hr timepoint, each column was treated with either: 50uM Doxorubicin (Fisher BP251610); or an equivalent dilution of Ethanol (Fisher BP2818100. For the 24hr timepoint, the same schema for treatment was used, but with half-doses. Each treatment contained eight biological replicates for each condition. After 18 hrs of incubation with each drug, cell viability, cytotoxicity, and Caspase-3/7 activity were measured using the ApoTox-Glo Triplex Assay (Promega) in a GloMax-Multi+ Reader (Promega). All data were normalized to the ethanol-treated scrambled siRNA control samples. siRNAs were designed to specifically-target the elephant *LIF6* gene. Sequences of the three LIF6-specific siRNAs used are as follows: 1) 5’-GAAUAUACCUGGAGGAAUGUU-3’, 2) 5’-GGAAGGAGGCCAUGAUGAAUU-3’, 3) 5’-CACAAUAAGACUAGGAUAUUU-3’ (Dharmacon). We also validated efficiency of the knockdown via qRT-PCR using the primer sets described earlier, which specifically the *LIF6* gene, and confirmed the combination of all three *LIF6* siRNAs was ~88%.

To determine if LIF6 was sufficient to induce apoptosis we synthesized and cloned (GeneScript) the African elephant LIF6 gene into the pcDNA3.1+C-DYK expression vector, which adds at DYK epitope tag immediately C-terminal to the LIF6 protein. We transiently transfected Chinese hamster ovary (CHO) cells or MEFs with LIF6_pcDNA3.1+C-DYK expression vector using Lipofectamine LTX according to manufacturer protocol and as described above, and assayed cell viability, cytotoxicity, and the induction of apoptosis using an ApoTox-Glo triplex assay. Mitochondrion membrane potential was assayed in CHO cells using the fluorometric Mitochondrion Membrane Potential Kit (Sigma MAK147) 48 hours after transfection.

### Evolutionary analyses of LIF genes

We used a Bayesian approach to date *LIF* duplication events implemented in BEAST (v1.8.3) (Rohland et al., 2010), including all identified African elephant, hyrax, and manatee *LIF* duplicates, as well as cannonical *LIF* genes from armadillo, sloth, aardvark, golden mole, and *LIF6* genes from Asian elephant, woolly and Columbian mammoth, straight-tusked elephant, and American Mastodon (Palkopoulou et al., 2018). We used the GTR substitution, which was chosen as the best model using HyPhy, empirical nucleotide frequencies (+F), a proportion of invariable sites estimated from the data (+I), four gamma distributed rate categories (+G) with the shape parameter estimated from the data, an uncorrelated random local clock to model substitution rate variation across lineages, a Yule speciation tree prior, uniform priors for the GTR substitution parameters, gamma shape parameter, proportion of invariant sites parameter, and nucleotide frequency parameter. We used an Unweighted Pair Group Arithmetic Mean (UPGMA) starting tree. The analysis was run for 10 million generations and sampled every 1000 generations with a burn-in of 1000 sampled trees; convergence was assessed using Tracer, which indicated convergence was reached rapidly (within 100,000 generations).

To constrain nodes we used normal priors with estimated confidence intervals, the root node was constrained to be 105 MYA, the root of Xenarthra was constrained to be 66 MYA, the root of Afrosoricida was constrained to be 70 MYA, the root of Afrosoricida-Macroselidea divergence constrained to be 75 MYA, the Elephantidea root was constrained to be 7.5 MYA, the Afrotheria root was constrained to be 83 MYA, the Paeungulata root was constrained to be 68 MYA, and the Proboscidea root was constrained to be 16 MYA. Divergence dates were obtained from www.timetree.org using the ‘Expert Result’ divergence dates.

We used the RELAX method to (Wertheim et al., 2015) test if duplicate LIF genes experienced a relaxation of the intensity of selection using the DataMonkey web server (Delport et al., 2010). The alignment included all duplicate LIF genes identified in the African elephant, hyrax, and manatee genomes, as well as cannonical *LIF* genes from armadillo, sloth, aardvark, golden mole, and *LIF6* genes from Asian elephant, woolly and Columbian mammoth, straight-tusked elephant, and American Mastodon. Alignment confidence was assessed using GUIDANCE2 (Sela et al., 2015) with the MAFFT (Katoh et al., 2005) algorithm and 100 bootstrap replicates.

## Acknowledgements

The authors would like to thank E. Palkopoulou and D. Reich (Harvard Medical School), the Broad Institute, M. Hofreiter (University of Potsdam) and H. Poinar (McMaster University) for providing access to the Proboscidean genomes, and U.M. Moll for providing the Bax/Bak KO MEF cell line.

## Author contributions

Writing - Original Draft: V.J.L and J.M.V; Writing - Review & Editing: V.J.L and J.M.V; Conceptualization: V.J.L and J.M.V; Investigation: J.M.M, M.S., S.C., and V.J.L.; Methodology: J.M.M, M.S.; Formal Analysis: V.J.L and J.M.V; Supervision and Project Administration: V.J.L.

---

**Figure S1.**
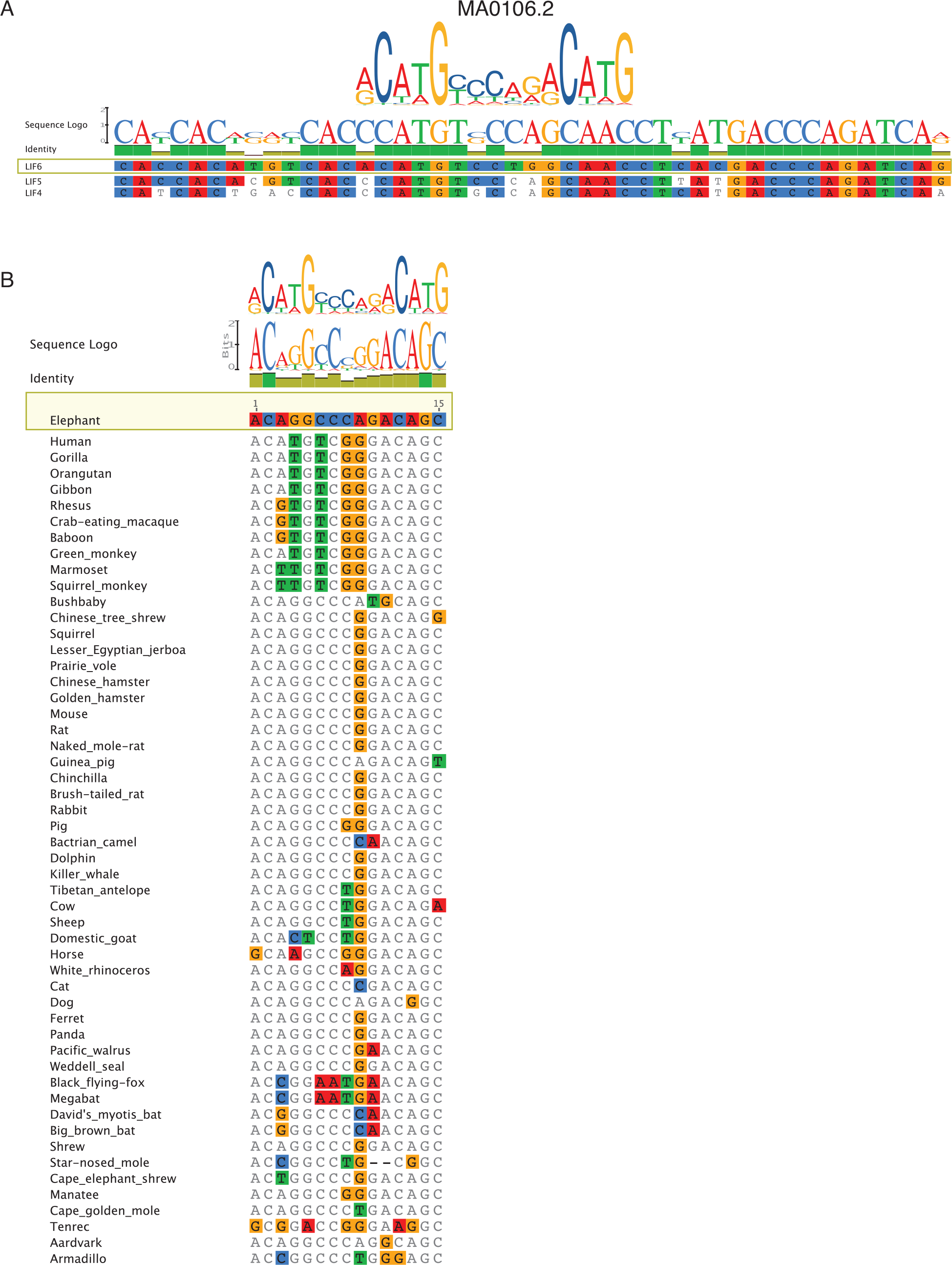
Similarity of the LIF6 and LIF1 TP53 binding sites to the TP53 binding motif (JASPAR MA0106.2). (A) Alignment of the LIF4, LIF5, and LIF6 TP53 binding sites. Bases are colored according to identity to LIF6, identical nucleotides are indicated with green columns above the alignment. A sequence logo is displayed on top. The experimentally validated TP53 binding motif is aligned on top of the putative LIF4, LIF5, and LIF6 TP53 binding sites. Note 3-4 nucleotide differences between LIF6 and LIF4 and LIF5. (B) Sequence logo of the LIF1 intron 1 TP53 binding site from 53 Eutherian mammals. The JASPAR TP53 motif (MA0106.2) is shown aligned and above a sequence logo of the TP53 motif from 53 mammals. Sequences from each of the 53 mammals is show below, with differences from the elephant LIF1 intron 1 TP53 binding site shown in color.

**Figure S2.**
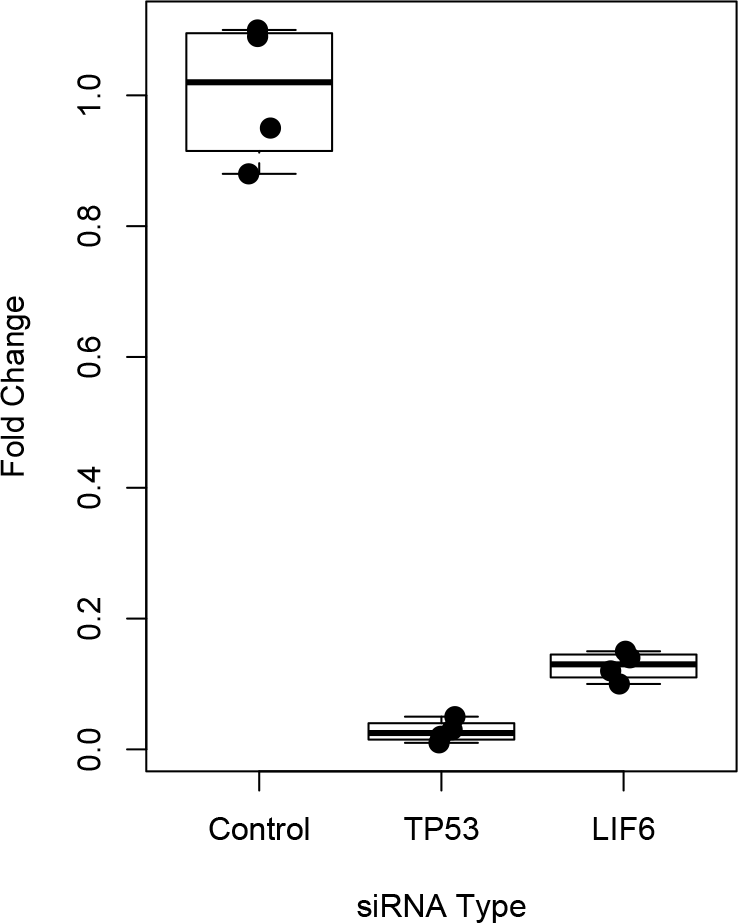
Efficacy of siRNAs targeting TP53 and LIF6 transcripts. Fold change in TP53 and LIF6 transcript abundance upon siRNA mediated knockdown compared to negative control siRNAs. N=4, Wilcox test *P*=0.028 for *TP53* knockdown and *P*=0.029 for *LIF6* knockdown.

**Figure S3.**
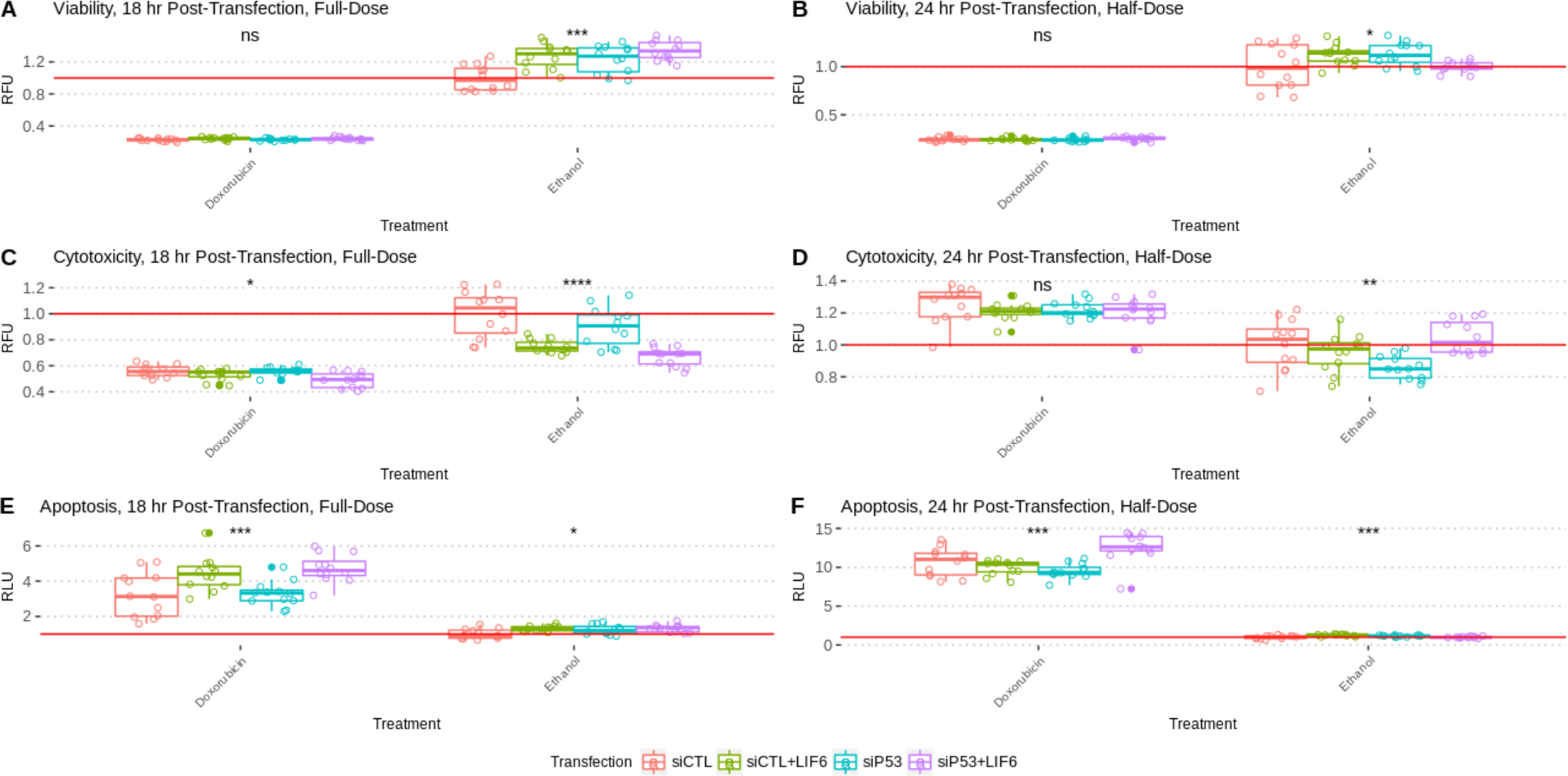
ApoTox-Glo results for elephant cells treated with LIF6 and siRNA to knockdown TP53, related to Figure 4E. Apoptosis (A,B), Cytotoxicity (C,D), and Viability (E,F) rates in African Elephant primary fibroblasts transfected with either scrambled control siRNA (siCTL) or anti-P53 siRNA (siP53); and with or without LIF6. After 6 hours of transfection, cells were treated with either 50-uM of Doxorubicin and tested 12 hours later at 18hr post transfection (A,C,E); or were treated with 25-uM Doxorubicin and tested 18 hours later at 24hr posttransfection (B,D,F). Co-transfecting siCTL with LIF6 results replicates the previously-seen apoptosis effect at 18 and 24 hours; at 24-hours, knocking down P53 rescues the apoptosis phenotype.

**Figure S4.**
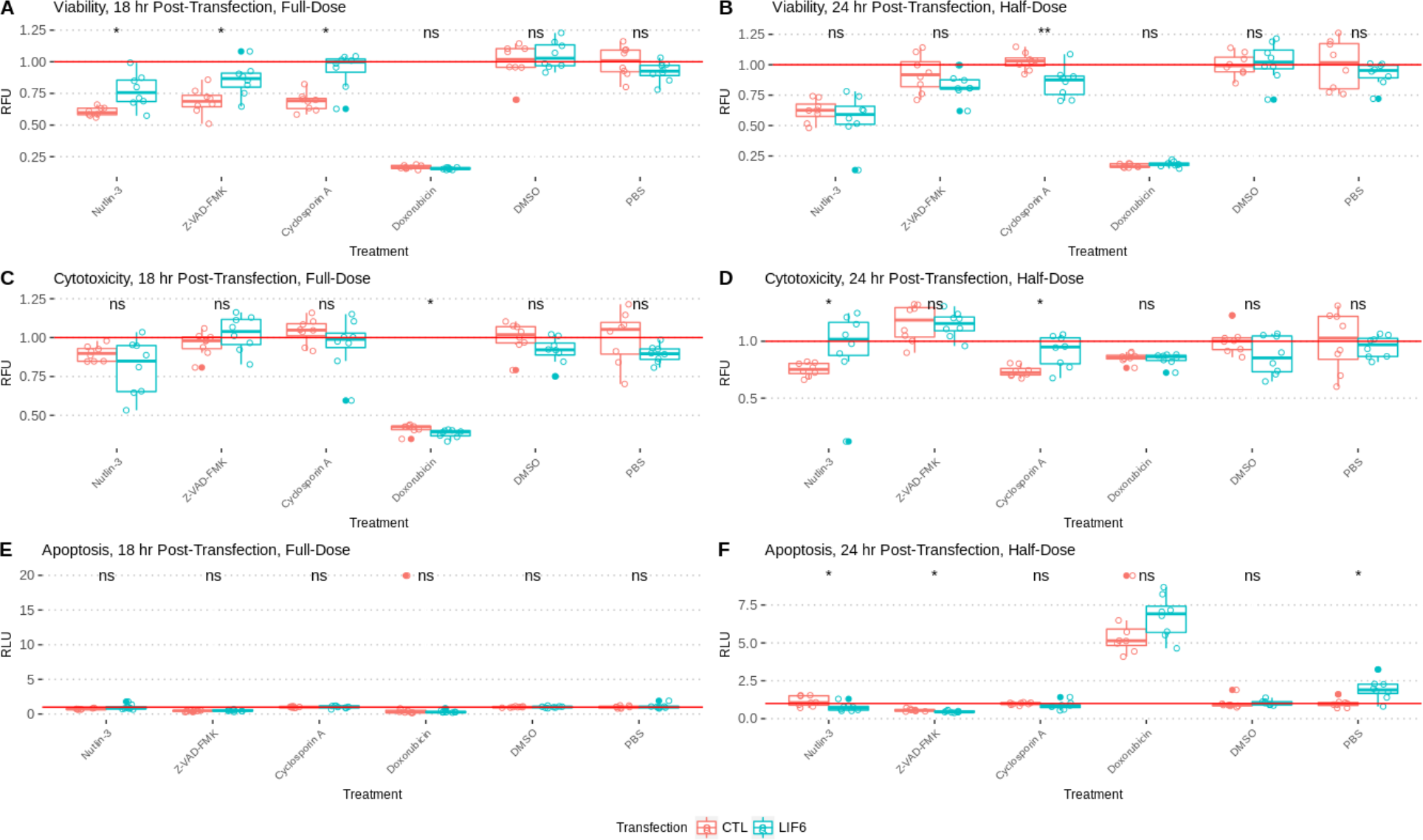
ApoTox-Glo results for elephant cells transfected with LIF6, related to Figure 5B. Apoptosis (A,B), Cytotoxicity (C,D), and Viability (E,F) rates in African Elephant primary fibroblasts transfected with LIF6, assayed the ApoToxGlo Triplex Assay. (A,C,E) Cells were treated 6 hours post-transfection with either 50-uM Nutlin-3, 20-uM Z-VAD-FMK, 2-uM Cyclosporin A, or 50-uM Doxorubicin, and were assayed 12 hours later, at 18 hours posttransfection. (B, D, F) Cells were treated as in A, C, and E, with half-doses of treatments, and tested 18 hours later at 24hr post-transfection. Apoptosis rates are markedly increased in cells transfected with LIF6 at 24 hours, which is inhibited by Z-VAD-FMK. Nutlin-3, which disrupts P53-MDM2 binding and thus activates P53, results in a increase in cytotoxicity, yet a decrease of apoptosis, in LIF6(+) cells compared to the mock-transfected control and to the PBS-treated LIF6(+) cells.

**Figure S5.**
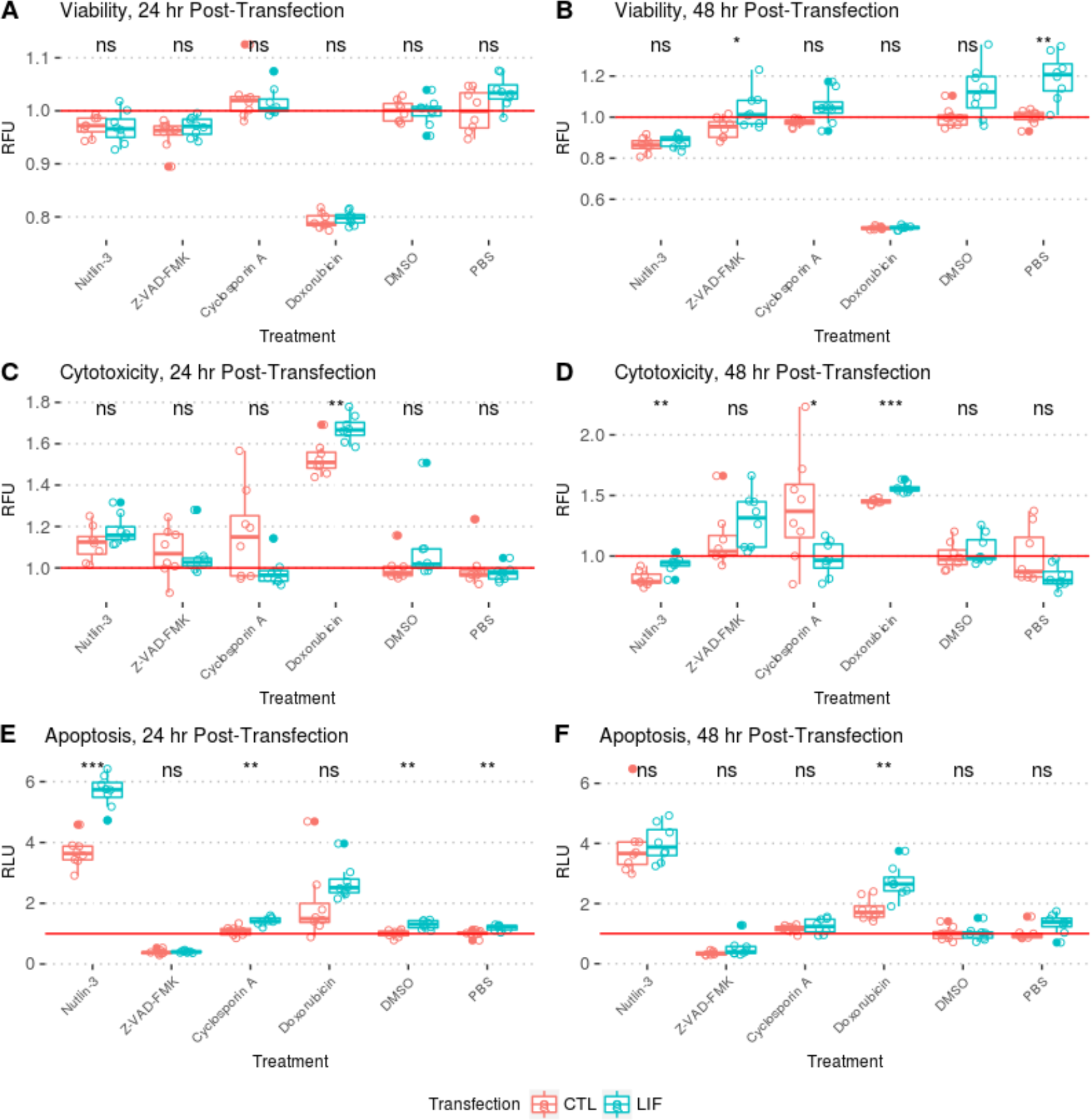
ApoTox-Glo results for mouse embryonic fibroblasts (MEFs) transfected with LIF6, related to Figure 6C. Cells were transfected with LIF6 for either 24-hours (A-C) or 48-hours (D-F), and were treated for 18 hours with either Nutlin-3, Z-VAD-FMK, Cyclosporin A, or Doxorubicin. The Viability (A, D), Cytotoxicity (B, E), and Apoptosis (C, F) rates in these cells were then measured using the ApoToxGlo Triplex Assay. Apoptosis rates are elevated for WT-MEF cells transfected with LIF6, but the effect is ablated when cells are treated with Z-VAD-FMK, a pan-caspase inhibitor; this ablation is not observed when treating cells with Cyclosporin A, an inhibitor of necrosis, indicating that the mechanism of LIF6-induced apoptosis is caspase-dependent. Treatment with Nutlin-3 - which increases P53 activity by disrupting binding between P53-MDM2 - intensifies apoptosis in LIF6 cells more than it does in untransfected WT-MEFs, suggesting a P53-dependent mechanism for caspase induction.

